# Analytical Solutions for the Time-Dependent Dynamics of Stochastic Gene Expression with mRNA-sRNA Interactions

**DOI:** 10.1101/2025.11.13.688103

**Authors:** Zhenhua Yu, Yiling Wang, Zhenyu Wang, Zhanpeng Shu, Zhixing Cao

## Abstract

The antagonistic interaction between small RNAs (sRNAs) and messenger RNAs (mRNAs) constitutes a fundamental regulatory mechanism of gene expression in both prokaryotic and eukaryotic cells. However, the stochastic nature of transcription renders mean-field approximations inadequate for quantitative analysis of such systems. In the regime of strong sRNA-mRNA antagonism, we generalize the conventional probability-generating-function (PGF) framework and derive a novel approximate solution in the form of a generalized PGF, which can be analytically transformed into the time-dependent joint distribution of sRNA and mRNA via Laurent series expansion. The proposed approximation accurately captures the full stochastic dynamics across diverse systems exhibiting strong antagonism, while incorporating key biological features such as transcriptional burstiness, translation and sRNA recycling over the entire temporal range. Building on this analytical foundation, we further develop a generalized-PGF-based parameter-inference method that enables efficient and precise estimation of kinetic parameters, achieving inference speeds up to three orders of magnitude faster than traditional maximum-likelihood estimation approaches.

## I. INTRODUCTION

Gene expression in living cells is orchestrated by a multilayered network of regulatory mechanisms. Beyond transcription factors and protein-protein interactions, a large class of non-coding RNAs, collectively termed small RNAs (sRNAs), plays a pivotal role in modulating gene expression at the post-transcriptional level. These molecules typically ranging from 20 to 300 nucleotides in length are widely found in both eukaryotes^1,2^ and prokaryotes^3,4^, where they participate in fine-tuning diverse physiological processes. In bacteria, sRNAs regulate key adaptive pathways in response to a variety of stresses, including oxidative^5^, osmotic^6,7^, acidic^8,9^, DNA damage^10^, and glucose-phosphate stress^11^, as well as processes such as quorum sensing^12^ and other regulatory circuits^13^. sRNAs also modulate virulence gene expression during host infection^3,14–16^, and play crucial roles in developmental canalization in mammalian cells^17,18^. Furthermore, dysregulation of sRNA biogenesis or activity has been closely associated with a variety of human diseases, including cancer^19–21^, highlighting their essential role in maintaining cellular homeostasis.

Among the various mechanistic modes of sRNA regulation, one of the most important and extensively studied is the antagonistic sRNA-mRNA interaction. In this mode, an sRNA molecule binds to its cognate mRNA target, often with the assistance of chaperone proteins, forming a duplex that suppresses translation and/or promotes degradation of one or both strands. Because of its central role in post-transcriptional control, understanding the kinetic and quantitative behaviour of antagonistic sRNA regulation is of great significance. In this stoichiometric regime, each sRNA molecule neutralizes a single mRNA molecule (or vice versa), as illustrated by Model I in Fig. 1. The dynamics of this process can be described by the following system of rate equations^22,23^,

**FIG. 1.**
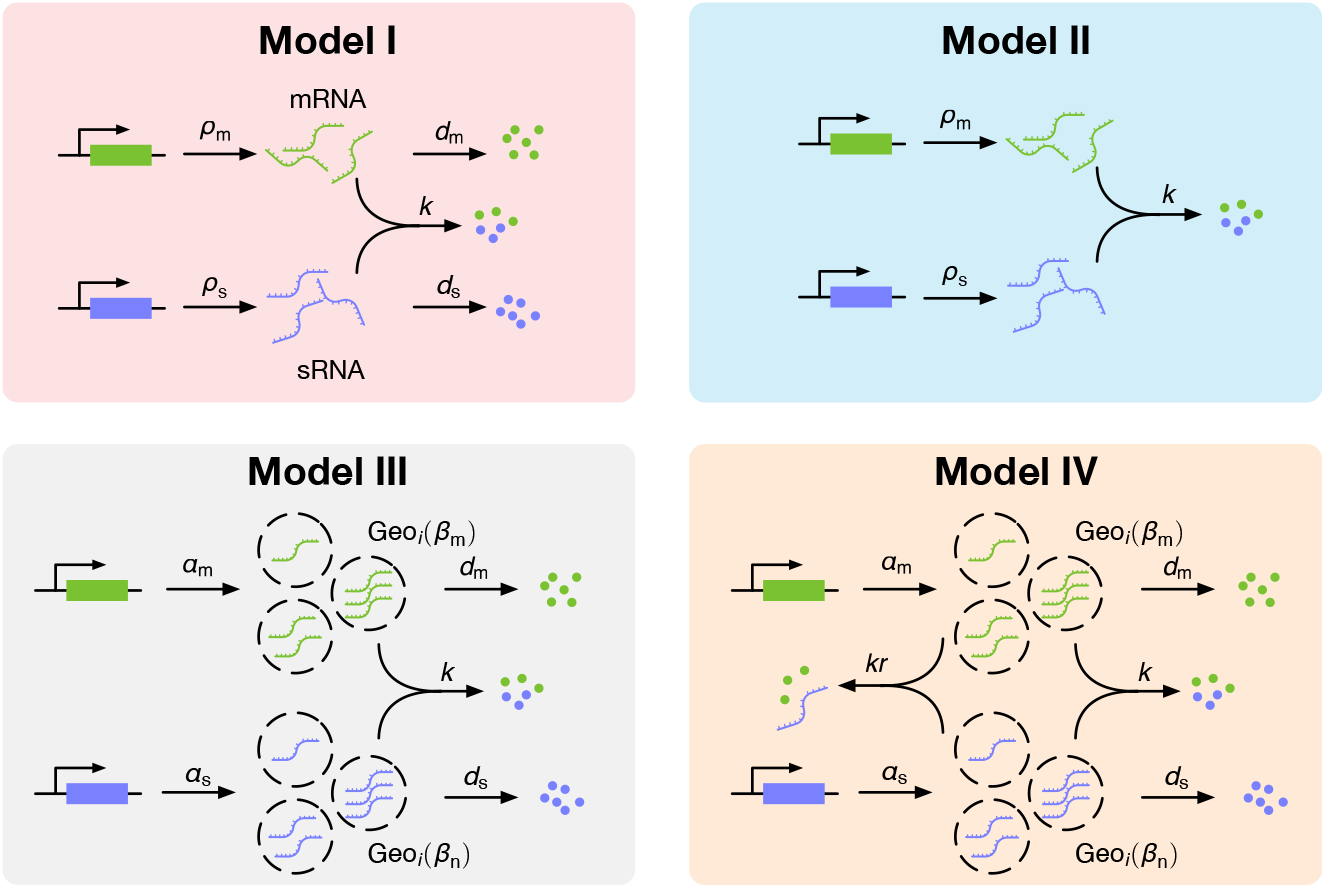
Schematic illustration of the four sRNA-mRNA interaction models analyzed in this study. Model I is the canonical antagonistic regulation model in which mRNA and sRNA are transcribed constitutively, degraded via first-order reactions, and mutually annihilated through a second-order binding reaction. Model II removes degradation of both mRNA and sRNA, retaining only constitutive transcription and mutual annihilation. Model III incorporates transcriptional bursting for both species, where each burst size follows a geometric distribution. Model IV extends Model III by including sRNA recycling, in which sRNA molecules are reused after repression and can neutralize multiple mRNAs during their lifetime.

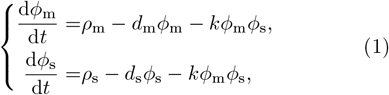

where *ϕ*_m_ and ϕ_s_ denote the cellular concentrations of mRNA and sRNA, *ρ*_m_ and *ρ*_s_ are their respective transcription rates, *d*_m_ and *d*_s_ are degradation rates, and *k* represents the binding rate constant. Shown in Refs.^22,23^, the steady-state solution to Eq. (1) exhibits a threshold-like response: when the transcription rate of the target mRNA is below a critical level, sRNAs fully repress expression; once the rate exceeds this threshold, residual mRNA escapes repression, producing a linear output. This characteristic nonlinearity underlies the switch-like behavior commonly observed in sRNA-mediated gene regulation.

While the deterministic rate-equation model in Eq. (1) captures the mean behaviour of antagonistic sRNA-mRNA regulation, it neglects the stochastic nature of gene expression^24–30^. Intrinsic noise arising from the probabilistic synthesis and degradation of RNA molecules can lead to substantial cell-to-cell variability in the degree of repression^31^. Moreover, antagonistic RNA interactions themselves play a crucial role in noise regulation, acting either to buffer or to amplify transcriptional fluctuations depending on the kinetic context^32–34^. Therefore, incorporating stochasticity is essential for accurately characterizing the kinetics and regulatory consequences of sRNA-mRNA antagonism. Nevertheless, it is mathematically challenging to analyse the stochastic kinetics because the antagonistic interaction introduces higher-order (nonlinear) propensities, leading to an *unclosed* hierarchy of moment equations. For instance, the first-order moment equations corresponding to Model I in Fig. 1 are

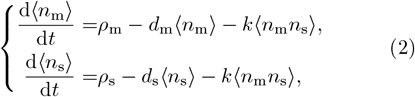

where ⟨•⟩ denotes the expectation operator, and *n*_*m*_ and *n*_*s*_ represent the numbers of mRNA and sRNA molecules, respectively. Eq. (2) reduces to the deterministic rate equations in Eq. (1) in the weak-antagonism limit, i.e., when the binding rate *k* is sufficiently small, which can be studied using perturbation analysis. However, this regime is biologically uninteresting, as weak antagonism implies negligible regulatory impact. Alternatively, one may employ moment-closure approximations to study the stochastic dynamics of sRNA and mRNA, such as the linear noise approximation (LNA)^35^, the normal moment closure^36,37^, or the linear mapping approximation (LMA)^38^. Nonetheless, the applicability assumptions underlying these methods often fail in this context. For example, the copy numbers of sRNA and mRNA can be extremely low, sometimes only a few molecules per cell, invalidating the continuum assumptions of the LNA. Similarly, the discrete and non-binary copy-number distributions of sRNA and mRNA challenge the foundational assumptions of the LMA, while the normal moment closure can yield non-physical or negative solutions^39,40^. These limitations were partially addressed in recent work^41^, which considered the strong antagonism regime – a biologically relevant condition where sRNA-mRNA binding occurs much faster than transcription or degradation. Under this assumption, the joint probability distribution of sRNA and mRNA copy numbers can be effectively reduced from a two-dimensional matrix to a one-dimensional vector, greatly simplifying the stochastic dynamics (detailed in Section II). This dimensional reduction enables an analytical expression for the steady-state distribution in the form of a probability-generating function (PGF), providing valuable mechanistic insight into the interplay between sRNA production, degradation, and target repression. Nevertheless, while many studies^41,42^ focus on steady-state behaviours, obtaining analytical results for time-dependent joint distributions of both sRNA and mRNA is far less common in the literature.

In this work, we develop an analytical framework for the time-dependent joint probability distribution of sRNA and mRNA molecules in the strong antagonistic regulation regime. By generalizing the PGF formalism, we obtain a closed-form approximate solution, the generalized PGF, which can be transformed into the full time-dependent joint distribution through Laurent series expansion. This formulation accurately captures stochastic dynamics over the entire timescale, accounting for biologically relevant features such as transcriptional burstiness and sRNA recycling. Our contributions are twofold: (i) we derive a tractable analytical solution for the time-dependent joint distribution of sRNA and mRNA under strong antagonism, expressed in the form of a generalized PGF; and (ii) we develop a generalized-PGF-based parameter inference method, which enables accurate estimation of kinetic parameters from time-series or snapshot data, achieving inference speeds up to three orders of magnitude faster than conventional maximum-likelihood approaches.

By bringing together stochastic chemical-kinetic modelling, analytical time-dependent distributions and parameter-inference methodology, our approach complements and extends previous work on sRNA-mediated regulation. It opens a path not only to deeper mechanistic insight into sRNA-mRNA kinetics but also to practical estimation of kinetics in experimental systems. In the following sections we will summarise the modelling approach, present solution derivation, and then apply the inference method to synthetic and (where available) experimental-style data. The remainder of this paper is organized as follows. Section II presents the stochastic models and introduces the strong-antagonism approximation. Section III motivates the use of a generalized probability generating function (GPGF). Sections IV to VI apply the GPGF approximation to derive time-dependent joint distributions for a range of mRNA-sRNA interaction models. Section VII demonstrates that the GPGF-based solutions significantly enhance the computational efficiency of parameter inference without sacrificing accuracy. Finally, Section VIII summarizes the key findings and discusses potential extensions.

## II. MODELS AND ASSUMPTIONS

In Fig. 1, we illustrate the four models considered in this study. *Model I* represents the canonical mRNA-sRNA antagonism model, in which both mRNA and sRNA are transcribed constitutively, degraded via first-order reactions, and mutually annihilated through a second-order interaction. This model has been extensively studied in the literature^22,23^. *Model II* simplifies the system by neglecting the degradation of mRNA and sRNA, while retaining the same transcription and annihilation mechanisms. *Model III* introduces a biologically more realistic feature – transcriptional bursting – for both mRNA and sRNA^27,28^. Here, transcription occurs intermittently, with long periods of transcriptional silence interspersed with brief, high-intensity bursts^43^. This process is modeled as:

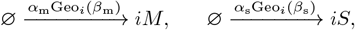

where the burst size parameters *β*_m_ and *β*_s_ follow a geometric distribution, defined as Geo_*i*_(*x*) = *x*^*i*^*/*(1 + *x*)^*i*+1^. *Model IV* extends Model III by incorporating sRNA recycling – an experimentally observed phenomenon in which sRNA molecules are reused after duplex-mediated repression^44^. This allows a single sRNA to neutralize multiple mRNAs over its lifetime. The recycling mechanism is modeled by the following second-order reaction

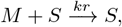

where *r* denotes the relative reaction rate coefficient with respect to the irreversible annihilation reaction

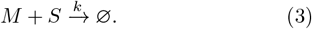

The stochastic dynamics of Model I in Fig. 1 are governed by the following chemical master equation (CME)

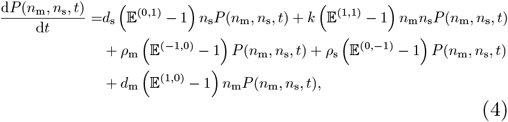

where *P* (*n*_m_, *n*_s_, *t*) denotes the probability of observing *n*_m_ mRNAs and *n*_s_ sRNAs in a cell at time *t*. The step operator 𝔼 (*i,j*) acts on a general function *f* (*n*_m_, *n*_s_) as 𝔼 (*i,j*)*f* (*n*_m_, *n*_s_) = *f* (*n*_m_ + *i, n*_s_ + *j*). For notational convenience, the time variable *t* is omitted hereafter. In general, solving Eq. (4) analytically is intractable, and one must typically resort to numerical methods such as the stochastic simulation algorithm (SSA)^45^ or the finite state projection (FSP) method^46^. However, reliance on such numerical approaches presents two key limitations: (i) they are computationally intensive, making them costly for parameter inference from large experimental datasets; and (ii) they offer limited analytical tractability, making it difficult to extract mechanistic or biological insights from the resulting solutions.

To obtain an analytical solution, one may consider the limit in which the binding rate constant *k* in Eq. (3) is small. However, this regime is of limited biological and physical interest, as the stochastic dynamics of the two interacting species are then well-approximated by independent Poisson processes, yielding trivial behavior with minimal coupling. Instead, following the approach in Ref.^41^, we focus on the opposite limit where *k* is large. Intuitively, under strong antagonism, mRNA and sRNA molecules cannot coexist in appreciable numbers: as soon as both *n*_m_ and *n*_s_ are nonzero, the annihilation reaction in Eq. (3) is immediately triggered, driving the system toward a state in which either mRNA or sRNA is depleted. Consequently, the joint probability satisfies *P* (*n*_m_, *n*_s_) = 0 for all *n*_m_ ≥ 1 and *n*_s_ ≥ 1. This observation allows us to derive a reduced set of CMEs, governing the dynamics confined to the *n*_m_-axis (*n*_s_ = 0), the *n*_s_-axis (*n*_m_ = 0), and the origin (*n*_m_ = 0, *n*_s_ = 0):

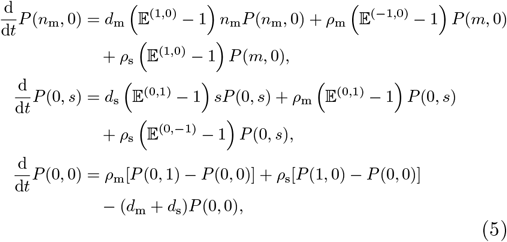

as previously derived in Ref.^41^. For completeness, a more rigorous derivation is provided in Supplementary Note 1. Under steady-state conditions, Eq. (5) can be solved analytically using the PGF method. Defining the PGF as

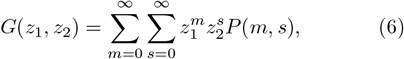

the steady-state solution in PGF form is given by

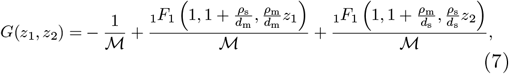

where 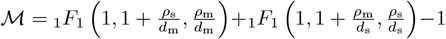. Here _1_*F*_1_ denotes the confluent hypergeometric function. The joint distribution of mRNA and sRNA can be constructed using the relation

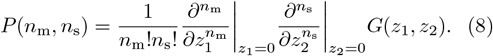

Direct evaluation of high-order symbolic derivatives with respect to two variables can be computationally intensive. A more practical alternative is to individually expand the second and third terms in Eq. (7) as Taylor series using the `Mathematica` command `NSeries`. The solution in Eq. (7) was originally reported in Ref.^41^, and a more streamlined derivation is provided in Supplementary Note 2.

Nevertheless, the solution in Eq. (7) has three notable limitations: (i) The analytical procedure based on Eqs. (5)–(8) is not readily generalizable to other dynamical systems exhibiting strong antagonism, such as Model III. As seen in the third equation of Eq. (5), the probability *P* (0, 0) depends only on *P* (1, 0) and *P* (0, 1). However, in Model III, due to transcriptional bursting, *P* (0, 0) is expected to depend on *P* (*i*, 0) and *P* (0, *i*) for *i >* 1, introducing higher-order dependencies. This complexity makes it difficult to apply traditional PGF method to derive analytical solutions. (ii) The closed-form expression in Eq. (7) involves confluent hypergeometric functions, which can pose challenges for numerical evaluation and may lead to instability during Taylor series expansion procedures. (iii) The current approach is restricted to the steady-state regime and cannot be directly extended to time-dependent solutions.

## III. MOTIVATING EXAMPLE

To overcome the aforementioned limitations, we generalize the PGF framework and employ it to analytically solve for the joint distributions of mRNA and sRNA copy numbers across Models I-IV in Fig. 1. Before presenting the general solution, we first consider a simplified case – Model II – in which both mRNA and sRNA degradation reactions are excluded.

Let *n*_m_(*t*) and *n*_s_(*t*) denote the number of mRNA and sRNA molecules at time *t*, respectively. Under the strong antagonism limit (*k*→ ∞), we define a new random variable *n*(*t*) = *n*_m_(*t*) − *n*_s_(*t*) to compactly represent the system’s state. In this formulation, *n*(*t*) *>* 0 indicates the presence of mRNA only, while *n*(*t*) *<* 0 corresponds to the presence of sRNA only, since mutual annihilation precludes coexistence of the two species.

Studying the variable *n*(*t*) is equivalent to analyzing the cumulative number of transcribed mRNAs and sRNAs up to time *t*, denoted by 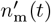 and 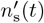 respectively. These quantities correspond to the number of molecules produced in two independent birth processes:

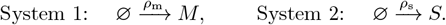

It follows directly that 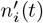 obeys a Poisson distribution, i.e., 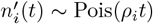 Pois(*ρ*_*i*_*t*) for *i* = *m* or *s*. The corresponding PGFs are

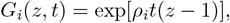

where *G*_*i*_(*z, t*) is defined in the conventional sense as

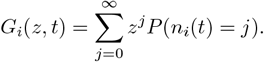

Alternatively, the PGF can be expressed in expectation form as

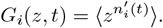

By the same principle, the PGF of *n*(*t*) can be written as

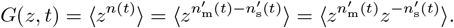

Since 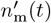 and 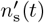 are independent, the PGF of *n*(*t*) factorizes as

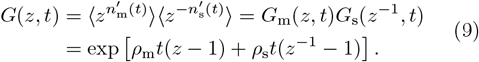

The expression above gives the PGF of *n*(*t*). However, unlike standard PGFs, which apply to non-negative integer-valued random variables, *n*(*t*) can take both positive and negative integer values. As such, its GPGF is naturally represented as a *Laurent series*

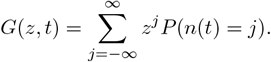

The coefficients of this Laurent series can be obtained analytically as

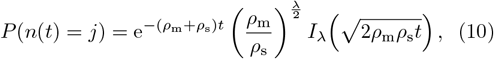

for any integer *λ*, where *I*_*λ*_(·) denotes the modified Bessel function of the first kind. The resulting probability distribution is known as the *Skellam distribution*, written as *n*(*t*) ~ Skel (*ρ*_m_*t, ρ*_s_*t*).

To validate the theoretical framework developed above, we simulated Model II in Fig. 1 using the Julia implementation of the SSA provided in Ref.^47^. A total of 10^6^ trajectories were generated up to time *t* = 2, starting from the initial condition with zero mRNA and sRNA molecules. Simulations were performed across a range of binding rate values *k*. In Fig. 2a, we compute the total probability mass confined to the “L”-shaped region of the state space at time *t* = 2, defined as

**FIG. 2.**
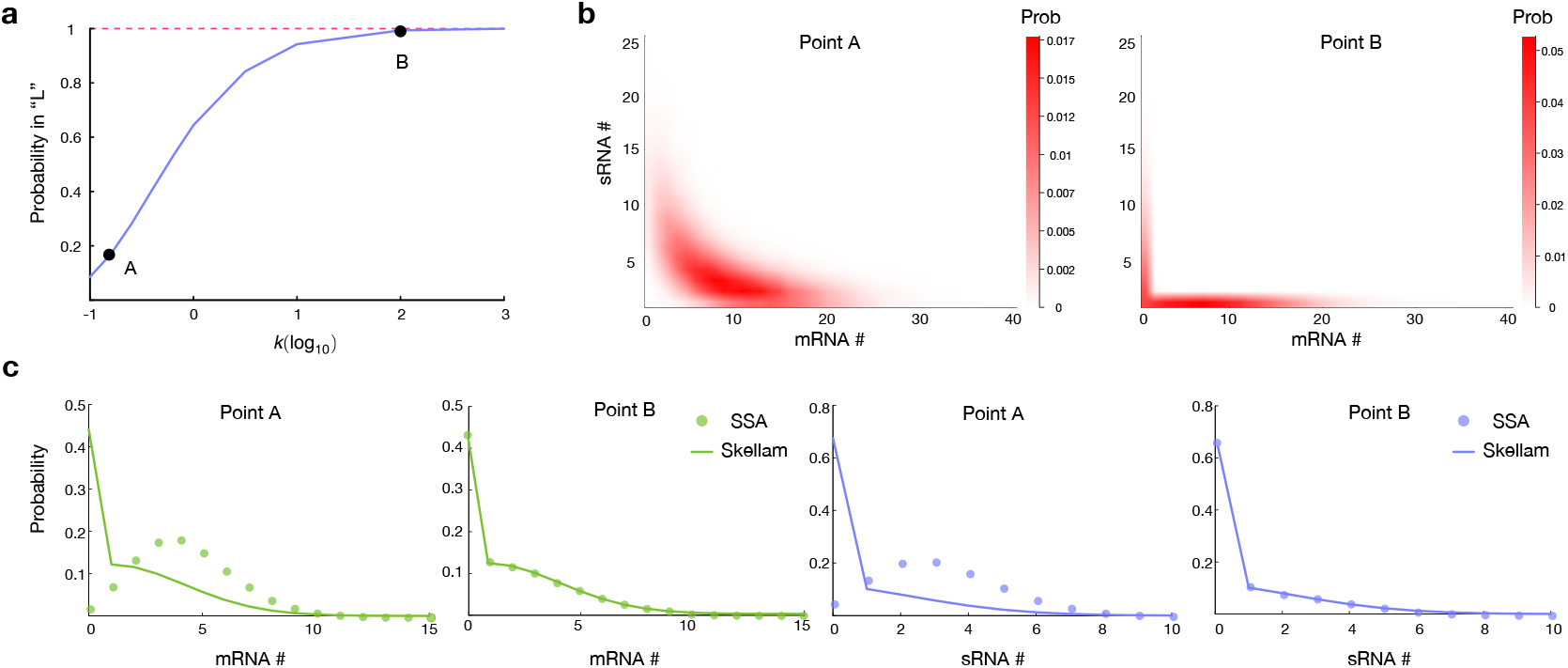
Validation of the strong antagonism approximation for Model II. (a) Total probability mass within the “L”-shaped region of the state space at time *t* = 2, computed across varying values of the binding rate constant *k*. As *k* increases, the probability outside this region approaches zero, confirming the theoretical prediction of mutual exclusivity between mRNA and sRNA. (b) Joint distributions of mRNA and sRNA copy numbers at two representative binding rates, corresponding to Point A (*k* = 10^0.8^) and Point B (*k* = 2) in panel (a). At Point B, the distribution exhibits a well-defined “L”-shape, characteristic of strong antagonism; at Point A, the structure is less pronounced but still reflects partial exclusion. (c) Marginal distributions of mRNA and sRNA obtained from SSA simulations (dots) compared with Skellam distribution predictions from Eq. (10) (solid lines) at Points A and B. Other simulation parameters: *ρ*_m_ = 3, *ρ*_s_ = 2.5, *t* = 2.

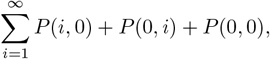

and observe that it approaches unity as *k* increases. This result confirms the theoretical prediction in Section II that, under strong antagonism, the probability outside this region vanishes. In Fig. 2b, we visualize the joint distributions of mRNA and sRNA at two representative binding rates, corresponding to Points A and B in Fig. 2a, where *k* = 10^0.8^ and *k* = 2, respectively. The joint distribution at Point A exhibits a partial “L”-shaped structure, while that at Point B shows a clear and well-defined “L” shape, consistent with the anticipated exclusion of states where both species coexist. In Fig. 2c, we compare the marginal distributions of mRNA and sRNA predicted by Eq. (10) at Points A and B with the ground-truth distributions obtained from SSA simulations. As the binding rate *k* increases, the theoretical predictions from Eq. (10) converge toward the SSA results, confirming the validity of the GPGF solution Eq. (9) (also Eq. (10)) in the strong antagonism regime.

The motivating example illustrates that the difference between two independent random variables offers a convenient and compact representation of their joint stochastic dynamics through a single composite variable. Furthermore, the GPGF given in Eq. (9) satisfies the following governing equation

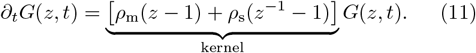

This observation highlights the role of the kernel in the time evolution of the GPGF and motivates its incorporation into a more general generating-function-based framework for analyzing Model I in Fig. 1.

## IV. REVISIT MODEL I

It is important to note that the key distinction between Model I and Model II lies in the presence of degradation reactions: Model I reaches a steady state due to the degradation of both mRNA and sRNA, whereas Model II does not. Motivated by the GPGF kernel in Eq. (11), we make a simplifying assumption that *d*_m_ = *d*_s_. Under this assumption, we heuristically propose the following generalized PGF equation to approximate the stochastic dynamics of mRNA and sRNA

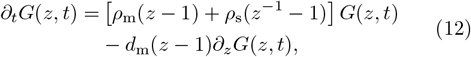

where the first term on the right-hand side corresponds to the transcription kernel defined in Eq. (11), and the second term accounts for the degradation of mRNA and sRNA, which now share the same rate *d*_m_.

By defining a new variable *u* = *z* −1, Eq. (12) can be rewritten as

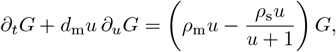

where, for simplicity, the explicit arguments of *G* are omitted without loss of clarity. Applying the method of characteristics yields the following system of ordinary differential equations (ODEs)

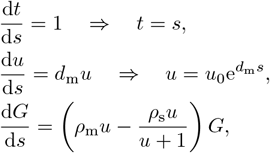

which admits the general solution

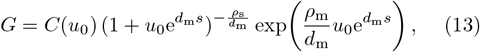

where *C*(*u*_0_) is a function of the initial condition.

Assuming the initial condition *g*(*u*) = *G* |_*t*=0_, we set *s* = 0 (equivalently, *t* = 0), yielding *u* = *u*_0_, *g*(*u*) = *g*(*u*_0_), and *G*|_*s*=0_ = *g*(*u*_0_). Substituting these relations gives

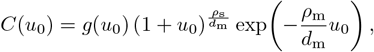

which can then be inserted into Eq. (13). Replacing 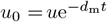, *s* = *t*, and *u* = *z* − 1 leads to the final solution:

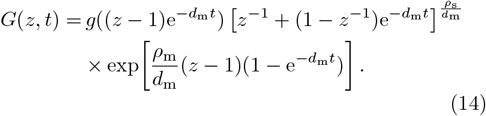

Given that *g*(0) = 1, the steady-state limit of Eq. (14) as *t*→ ∞ yields a remarkably simple expression for the GPGF

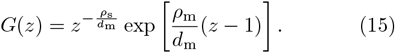

Surprisingly, both the time-dependent solution in Eq. (14) and the steady-state solution in Eq. (15) are significantly simpler than the exact PGF solution in Eq. (7), which involves confluent hypergeometric functions. Moreover, Eq. (15) reveals an intriguing structure: the joint distribution corresponds to a left-shifted Poisson distribution, where the prefactor 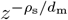 acts as a shift operator in the probability state space under the inverse *z*-transform.

Although the expressions in Eqs. (14) and (15) are remarkably simple, they rely on the strong assumption that *d*_m_ = *d*_s_. We next investigate the validity of this approximation when this constraint is relaxed, while still maintaining the strong antagonism condition (*k*→ ∞). To do so, we fix *ρ*_s_ = 10 and *d*_m_ = 1, and consider the steadystate regime. We vary *ρ*_m_ and *d*_s_ independently over an order of magnitude to assess how deviations from *d*_m_ = *d*_s_ influence the approximation. Since the GPGF is defined as a Laurent series, we numerically extract its coefficients using the `Mathematica` command `NSeries` to obtain the approximate joint distribution of mRNA and sRNA. For comparison, we compute the exact distribution by performing a Taylor series expansion of Eq. (7). The discrepancy between the approximate and exact distributions is quantified via the Wasserstein distance (WD), which we map as a function of the parameter ratios *ρ*_m_*/ρ*_s_ and *d*_s_*/d*_m_, as shown in Fig. 3a. Surprisingly, the approximation error decreases monotonically with increasing *ρ*_m_*/ρ*_s_, and remains largely insensitive to changes in *d*_s_*/d*_m_. This suggests that the relative transcription rate of mRNA versus sRNA is the dominant factor influencing the approximation accuracy, while the degradation rate *d*_s_ plays a negligible role when *ρ*_m_ *> ρ*_s_. In Fig. 3b, we illustrate the marginal distributions of mRNA and sRNA at two representative parameter settings (Points A and B in Fig. 3a). Point A corresponds to the largest WD and hence the poorest approximation, while Point B reflects a moderate case. Remarkably, even in the worst-case scenario (Point A), the approximate solution remains qualitatively consistent with the exact result; in the moderate case (Point B), the agreement is nearly perfect. The insensitivity to the degradation rate *d*_s_ is intuitive: since sRNA is less abundant than mRNA and primarily degraded via the interaction in Eq. (3), the effect of its intrinsic degradation is negligible. Due to the symmetry in Model I, when *ρ*_m_ *< ρ*_s_, one can reasonably expect that replacing *d*_m_ with *d*_s_ in Eq. (12) would yield a similarly accurate approximate GPGF solution.

**FIG. 3.**
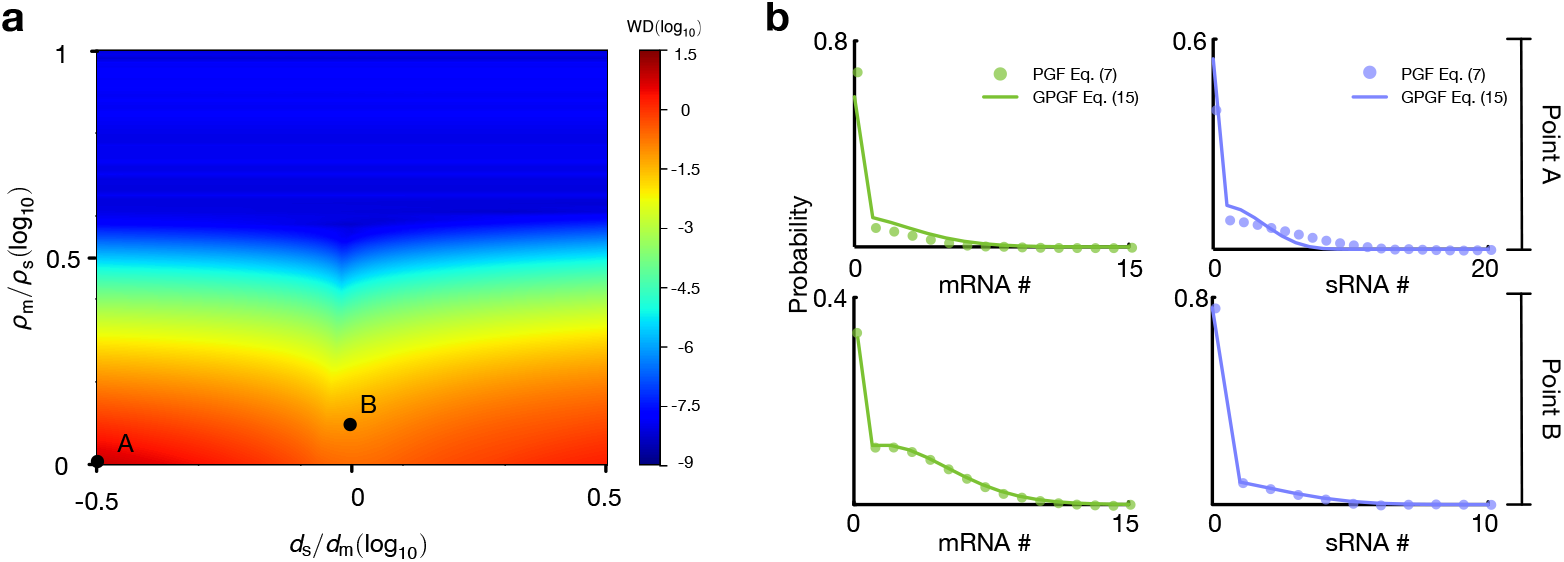
Quantifying the accuracy of the steady-state approximation under unequal degradation rates in Model I. (a) Wasserstein distance between the exact joint distribution (obtained from Eq. (7)) and the approximate distribution (derived from Eq. (15)) as a function of the transcription rate ratio *ρ*_m_*/ρ*_s_ and degradation rate ratio *d*_s_*/d*_m_, with *ρ*_s_ = 10 and *d*_m_ = 1 held fixed. (b) Marginal distributions of mRNA and sRNA copy numbers at two representative points from panel (a): Point A (worst-case approximation) and Point B (moderate approximation). Dots represent exact distributions computed from Eq. (7), and solid lines represent the approximate distributions based on Eq. (15). The remaining parameters are *ρ*_m_ = 10, *d*_s_ = 10^−0.5^ for Point A, and *ρ*_m_ = 10^0.08^, *d*_s_ = 1 for Point B.

In the top panel of Fig. 4, we demonstrate that the approximate marginal distributions of mRNA and sRNA derived from Eq. (14) are in excellent agreement with those obtained from SSA simulations (10^6^ trajectories) across multiple time points, under parameter conditions where the strong antagonism assumption remains valid. To the best of our knowledge, no analytical or approximate time-dependent solution has been previously reported for this regime.

**FIG. 4.**
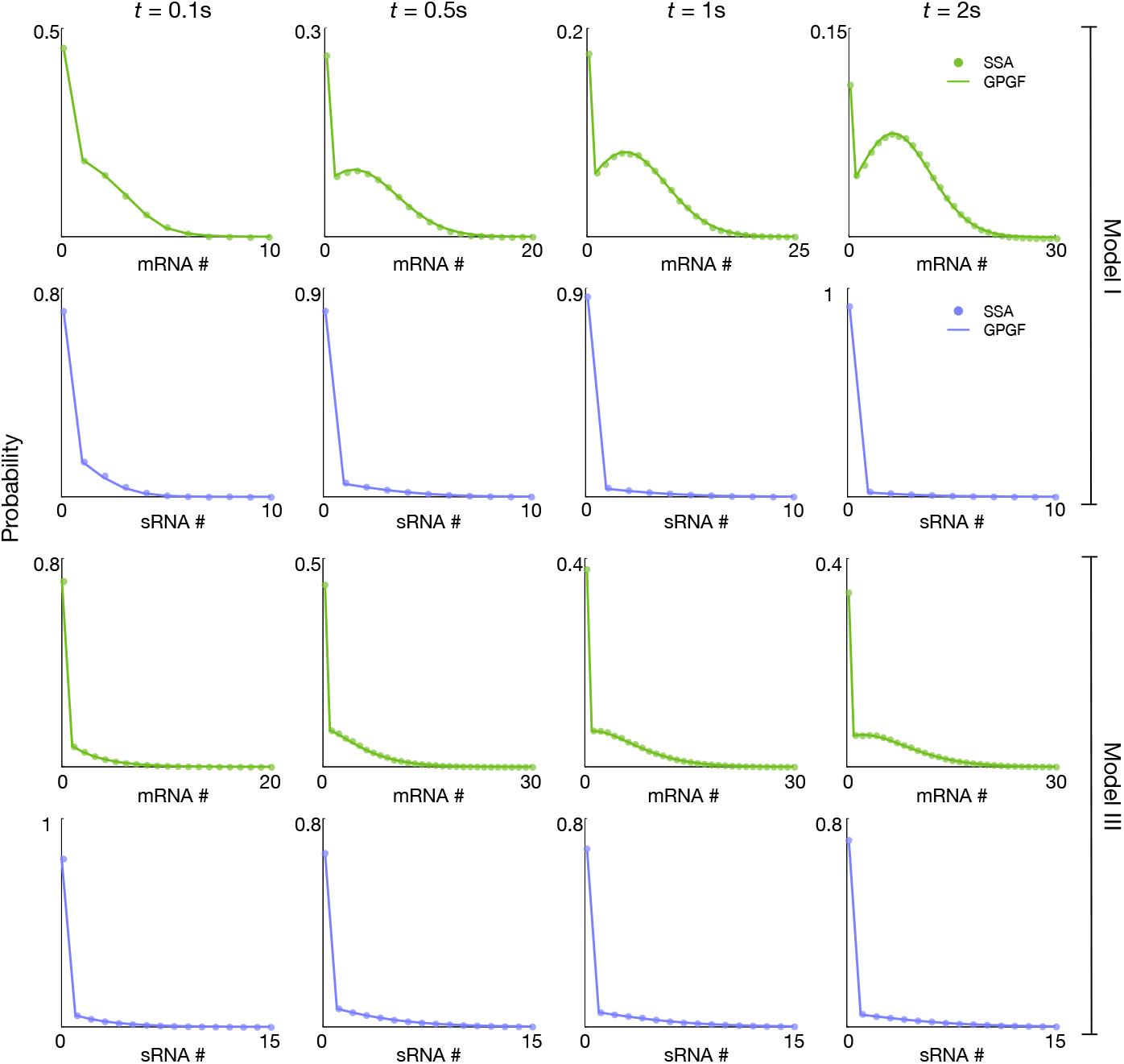
Validation of GPGF-based time-dependent approximations for Models I and III under strong antagonism (*k* = 10^4^). Top: Time evolution of marginal mRNA and sRNA distributions predicted by the analytical GPGF solution for Model I (Eq. (14), solid lines) shows excellent agreement with SSA simulations (dots, 10^6^ trajectories). Bottom: The GPGF-based marginal distributions for Model III (Eq. (18)) are likewise in close agreement with SSA results, confirming the validity of the approximation in the bursty transcription regime. Simulation parameters: Model I – *ρ*_m_ = 30, *ρ*_s_ = 22, *d*_m_ = *d*_s_ = 1; Model III –*α*_m_ = 6, *α*_s_ = 4, *β*_m_ = 2, *β*_s_ = 2, *d*_m_ = *d*_s_ = 1.

## V. MODEL III

The central step in the GPGF approximation is to formulate the governing equation – specifically, to derive the corresponding kernel. We now apply this GPGF-based framework to seek an analytical time-dependent solution for Model III, which is intractable using the traditional PGF method outlined in Section II. The key challenge arises from transcriptional burstiness, which permits a one-step jump from state *i* to −*j*: that is, a burst event may produce *i* + *j* sRNAs, neutralizing *i* mRNAs and leaving *j* sRNAs unbound – effectively bypassing the intermediate state 0. This non-local transition violates the structure that allowed analytical solution of the PGF equations in Eqs. (S5)–(S7) (see Appendix Supplementary Note 2). As a result, neither the steady-state nor the time-dependent PGF for Model III can be solved using the classical techniques used for Model I.

To analyze Model III, we adopt a strategy similar to that used for Model I by first considering a simplified variant in which degradation reactions are excluded

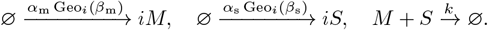

As before, we define a single random variable *n*(*t*) = *n*^*′*^_m_(*t*) − *n*^*′*^_s_(*t*) to capture the net copy number difference between mRNA and sRNA, under the strong antagonism condition (*k*→ ∞). The transcription processes *n*^*′*^_m_(*t*) and *n*^*′*^_s_(*t*) are independent and governed by

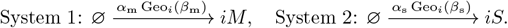

Each of these systems is a compound Poisson process with geometric jump size, yielding a *Pólya–Aeppli* distribution, i.e.,

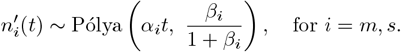

The PGF of such a process can be derived by replacing *z* with the PGF of geometric distribution [1 − *β*_*i*_(*z* − 1)]^−1^ in the standard birth-process PGF exp[*α*_*i*_*t*(*z* − 1)] (see Supplementary Note 2 of Ref.^48^):

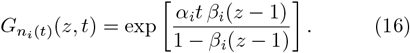

Thus, the GPGF for *n*(*t*) becomes

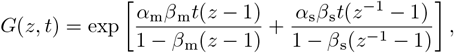

which satisfies the governing equation:

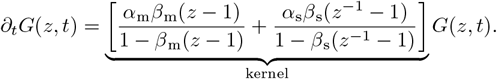

This expression provides the kernel needed for GPGF approximation for Model III. To approximate the full model, we now incorporate degradation terms as in Section III. Assuming mRNA is more abundant (*ρ*_m_ *> ρ*_s_), we introduce degradation of mRNA only, leading to the governing equation for Model III

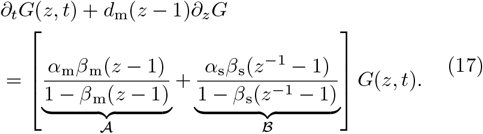

Assuming zero initial copy numbers of both mRNA and sRNA, we solve Eq. (17) using the method of characteristics. The resulting analytical time-dependent GPGF for Model III is:

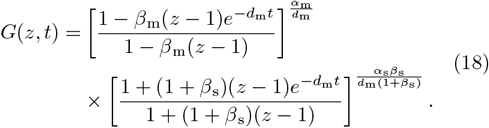

In the bottom panel of Fig. 4, we also show that the marginal distributions of mRNA and sRNA predicted by the analytical solution in Eq. (18) closely match those obtained from SSA simulations (10^6^ trajectories) across multiple time points, thus validating the accuracy of the GPGF approximation for Model III.

## VI. MODEL IV

Next we apply the GPGF approximation to analyze the distribution dynamics of Model IV in Fig. 1, where the interaction between mRNA and sRNA leads to two distinct outcomes: either degradation of the mRNA or co-degradation of both mRNA and sRNA. The reaction *M* + *S* →*S* captures the recycling behavior of sRNA, as reported in Refs.^23,44^. Under the strong antagonism assumption (*k* → ∞), the following reactions

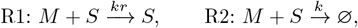

proceed at the fastest time scale. As such, each sRNA molecule can potentially degrade multiple mRNAs before itself being degraded. Specifically, at each reaction event between R1 and R2, the probability of R1 occurring is *r/*(1 + *r*), and that of R2 is 1*/*(1 + *r*). If an sRNA is degraded after exactly *i* −1 recycling events (i.e., *i* total mRNA degradations), the probability is

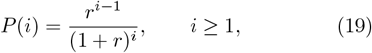

which follows a geometric distribution, and its corresponding PGF is

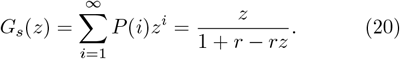

This motivates us to conceptualize each sRNA molecule as a package of virtual sRNAs (vsRNAs) (illustrated as the red curves in Figs. 5a and 5b), where each vsRNA degrades exactly one mRNA. The number of vsRNAs per sRNA follows the geometric distribution in Eq. (19). Therefore, the total number of vsRNAs at time *t* follows a compound distribution – a Pólya-Aeppli distribution (from sRNA production) composed with the geometric distribution of vsRNAs per sRNA. The PGF of the vsRNA count, denoted *n*_vs_(*t*), is obtained by substituting *z* in Eq. (16) with *G*_*s*_(*z*) from Eq. (20), yielding

**FIG. 5.**
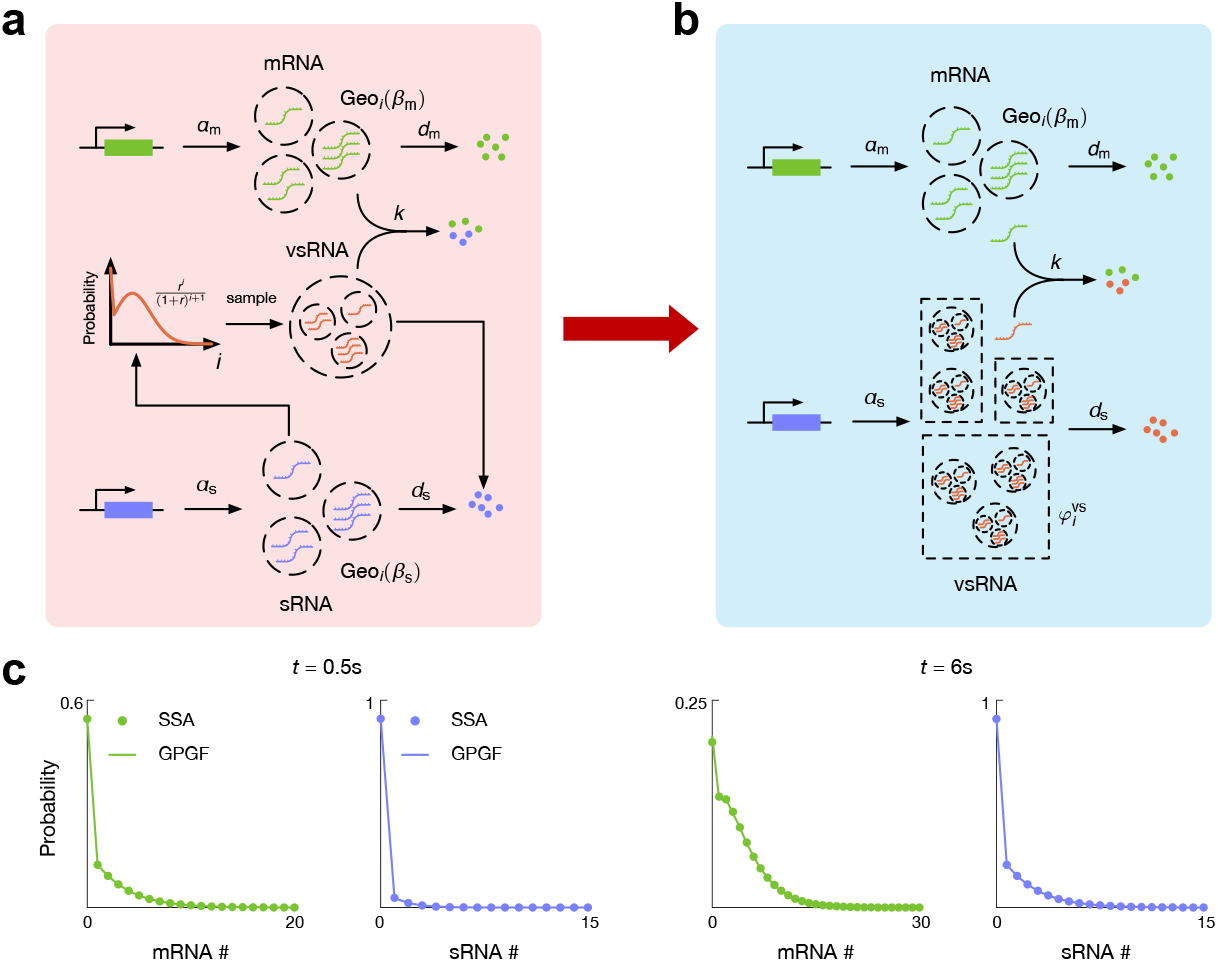
GPGF-based analytical approximation for Model IV under strong antagonism (*k*→ ∞). (a)(b) Illustration of the concept of virtual sRNA (vsRNA), where each vsRNA corresponds to one mRNA-degrading unit, and the total vsRNA count per sRNA follows a geometric distribution. (c) Time-dependent marginal distributions of mRNA (gree) and sRNA (blue) at *t* = 0.5 and *t* = 6, obtained from the GPGF-based analytical solutions (Eqs. (24) and (29), solid lines) and validated against SSA simulations (10^6^ trajectories, dots). The parameters used for simulations are *a*_m_ = 2.2, *a*_s_ = 0.7, *b*_m_ = 2, *b*_s_ = 1, *d*_m_ = 1, *d*_s_ = 1.2 and *k* = 10^4^.

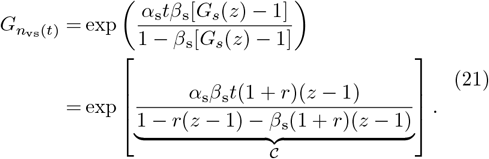

To derive the GPGF governing equation for Model IV, we follow the same procedure outlined in Eqs. (16)–(17). Specifically, we focus on the two transcription processes illustrated in Fig. 5b,

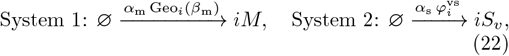

where the species *S*_*v*_ represents vsRNA, and 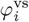 denotes the burst size distribution for vsRNA, defined as the compound distribution resulting from the distribution given in Eq. (19) and the geometric distribution Geo_*i*_(*β*_s_). The explicit form of 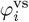 is not required for the subsequent derivation and will therefore not be computed. We also note that the PGF of System 2 in Eq. (22) has already been provided in Eq. (21). Then we substitute *z* with 1*/z* in the expression for in Eq. (21), and then replace in Eq. (17) with the resulting expression. This yields the GPGF equation for Model IV

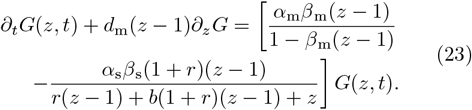

Assuming an initial condition of zero mRNA and sRNA, Eq. (23) can be solved (via the method of characteristics) to yield the analytical GPGF

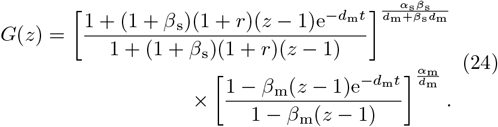

Importantly, when expanding *G*(*z*) in Eq. (24) as a Laurent series, the coefficients of *z*^*n*^ for *n >* 0 represent the probabilities of mRNA counts. However, the coefficients for *n <* 0 correspond to the probabilities of vsRNA counts rather than actual sRNA copy numbers.

Hence, we next derive the distribution of sRNA. To do so, we must convert the distribution of vsRNA back into that of sRNA, which amounts to a decompounding operation, i.e., removing the influence of the distribution defined in Eq. (19) from the burst size distributions of both Systems 1 and 2 in Eq. (22), namely Geo_*i*_(*β*_m_) and 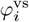. Once this is achieved, the transcription kernel governing the sRNA distribution can be determined from the two corresponding transcription processes,

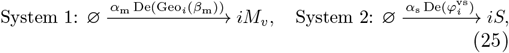

where De(•) denotes the decompounding operator, and *M*_*v*_ represents virtual mRNA, the counterpart to vsRNA. Decompounding 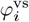 is straightforward, as it immediately yields Geo_*i*_(*β*_s_). To decompound Geo_*i*_(*β*_m_), we define an auxiliary variable *y* and solve for *z* from

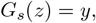

to obtain *z* as a function of *y*, which yields

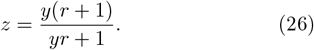

The decompounding of Geo_*i*_(*β*_m_) then amounts to substituting *z* in its PGF with Eq. (26), and finally replacing *y* back with *z*. However, this step can be bypassed by directly applying the same transformation to the PGF of System 1 in Eq. (22), i.e., applying it to term 𝒜 in Eq. (17). This gives the PGF for *M*_*v*_,

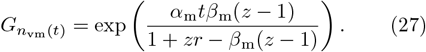

Accordingly, the GPGF governing equation for the sRNA distribution becomes

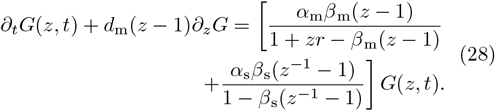

Solving Eq. (28) using the method of characteristics yields

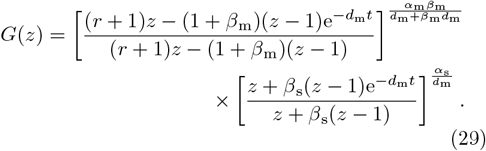

Expanding Eq. (29) in a Laurent series, the coefficients of *z*^*n*^ for *n <* 0 provide the distribution of sRNA counts. Taken together, the Laurent series expansions of Eqs. (24) and (29) yield the joint distribution of mRNA and sRNA under the strong antagonism assumption for Model IV.

We further compare the approximate marginal distributions of mRNA and sRNA derived from Eqs. (24) and (29) with those obtained via SSA simulations (10^6^ trajectories), as shown in Fig. 5c. The excellent agreement at both *t* = 0.5 and *t* = 6 provides strong validation for the GPGF-based approximation developed for Model IV.

## VII. PARAMETER INFERENCE BASED ON GPGF

Finally, we demonstrate how the GPGF approximate solution can enhance both the accuracy and efficiency of parameter inference. Specifically, we consider Model I in Fig. 1 under steady-state conditions. In this case, the binding rate *k* does not appear in the approximate steady-state solution (Eq. (15)), nor does the degradation rate of sRNA *d*_n_, since mRNA is assumed to be more abundant than sRNA. Likewise, the mRNA degradation rate cannot be inferred under the steady-state assumption. Consequently, only the two transcription rates, *ρ*_m_ and *ρ*_n_, need to be estimated, and forming the parameter vector *ϕ* = [*ρ*_m_, *ρ*_s_]^⊤^.

Following the framework presented in Ref.^49^, we consider *n*_*c*_ single cells, where the *i*-th cell contains *n*_m,*i*_ mRNAs and *n*_s,*i*_ sRNAs. The empirical GPGF (EGPGF) is then defined as

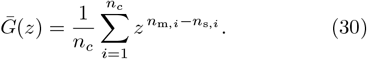

The parameters *ϕ* are inferred by minimizing the discrepancy between the model-predicted GPGF, *G*(*z, ϕ*), obtained from Eq. (15), and the empirical GPGF 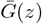. Specifically, the objective function is defined as

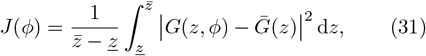

where 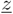 and 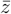 denote the lower and upper bounds of the integration, respectively. The loss function *J*(*ϕ*) thus represents the mean squared deviation between the model and empirical GPGFs over the specified interval.

To improve the computational efficiency of evaluating Eq. (31), we approximate the integral using Gauss–Legendre quadrature:

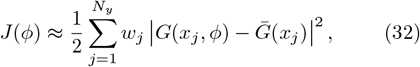

Where

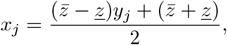

with *y*_*j*_ ∈ [−1, 1] denoting the *j*-th quadrature node of order *N*_*y*_, and *w*_*j*_ the corresponding weight, both obtained via the `gausslegendre` function in Julia.

Finally, we minimize *J*(*ϕ*) using the Nelder–Mead algorithm implemented in the `Optim.jl` package. This algorithm is selected for its robustness to initialization and its ability to avoid costly gradient evaluations of ∂_*ϕ*_*J*(*ϕ*). The complete numerical workflow is summarized in Algorithm 1 and illustrated in Fig. 6a. Unless otherwise stated, optimization is performed with a gradient tolerance of `g_tol` = 10^−30^ and a maximum of `iterations` = 1000, consistent with the default settings of the optimize function.

**FIG. 6.**
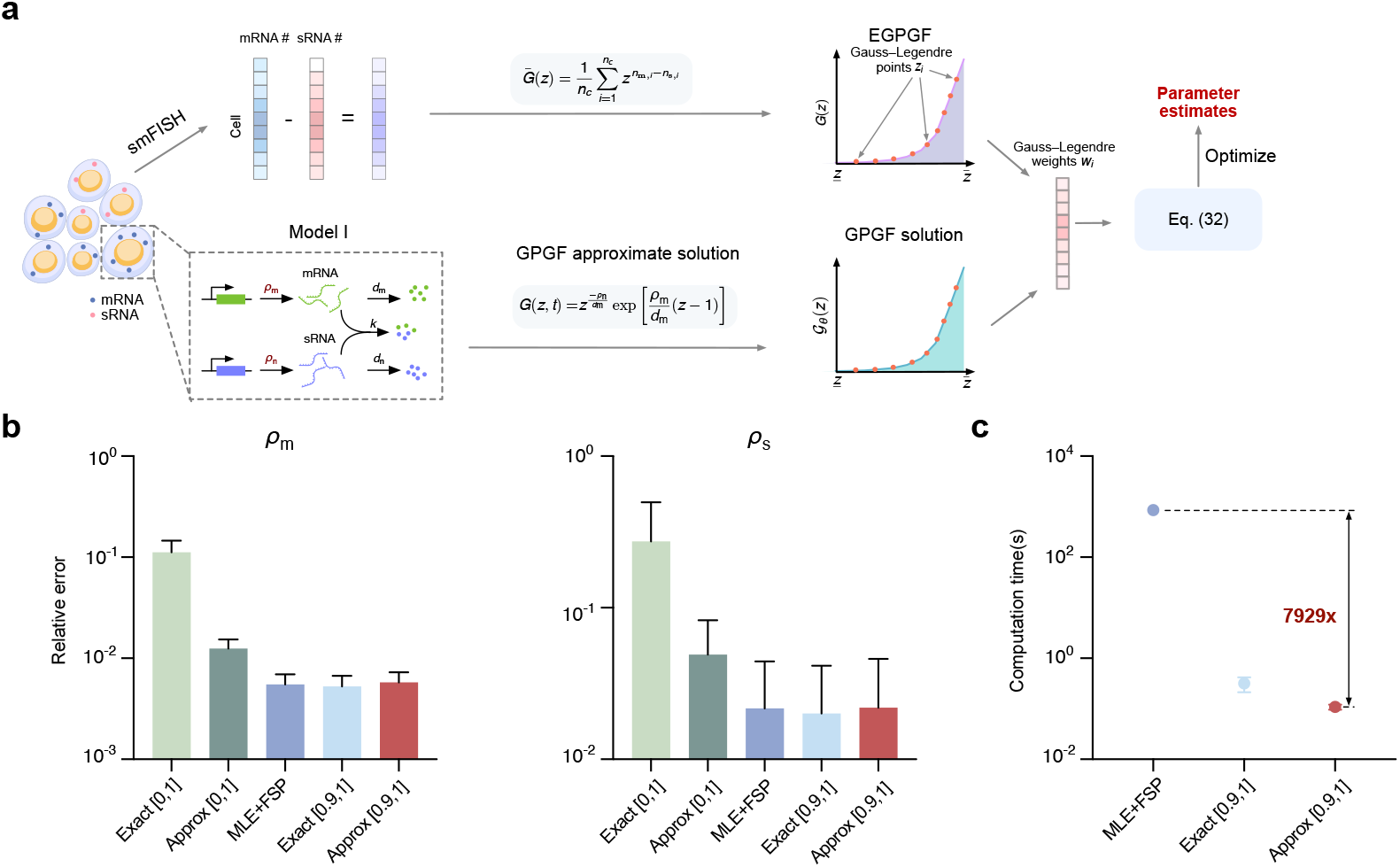
Performance comparison of different parameter inference protocols. (a) Schematic of the workflow for GPGF-based parameter inference using the approximate steady-state solution. (b) Relative inference errors of *ρ*_m_ and *ρ*_n_ across ten simulated datasets under five inference protocols: (i)–(ii) GPGF-based inference using the approximate solution Eq. (15), (iii)–(iv) PGF-based inference using the exact solution Eq. (7), and (v) MLE with the FSP method. (c) Comparison of computation times for the three best-performing protocols, showing that the GPGF-based method with integration interval [0.9, 1] (protocol (ii)) is the most efficient one.

We randomly sampled ten pairs of transcription rates (*ρ*_m_, *ρ*_n_) (Table S1), fixing *d*_m_ = *d*_n_ = 1 and *k* = 10^4^ to satisfy the strong antagonism condition. For each parameter pair, *n*_*c*_ = 10^4^ single-cell trajectories were simulated to steady state using SSA, corresponding to a typical sample size in single-cell sequencing. We compared five inference protocols: (i) GPGF-based inference using Eq. (15) with 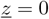 and 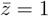(ii) the same with 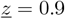 and 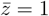; (iii) PGF-based inference (Ref.^49^, Supplementary Note 3) using the exact solution Eq. (7) with 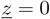 and 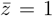; (iv) the same with 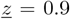 and 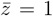; and (v) the maximum likelihood estimator (MLE) using the FSP method (Supplementary Note 3).

### Algorithm 1

GPGF-based inference method

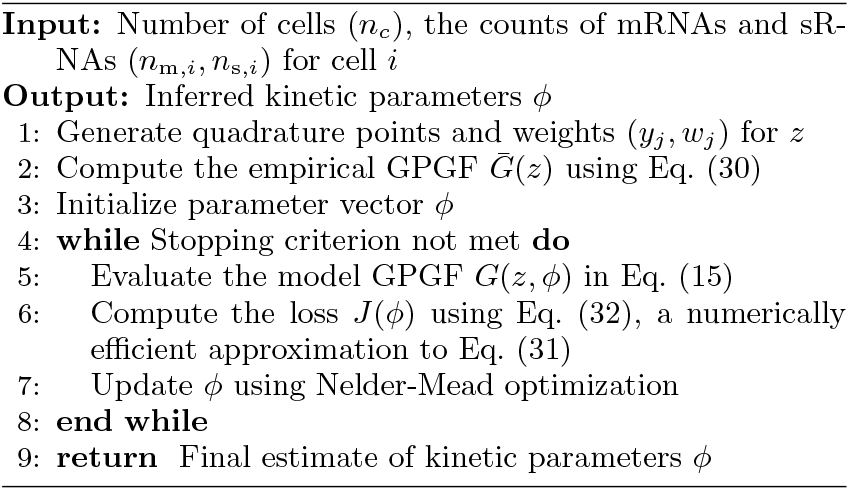

All inference tasks were executed on a MacBook Pro with an M2 chip and 32 GB memory. For each protocol, we calculated the relative errors of the inferred *ρ*_m_ and *ρ*_n_ across the ten simulated datasets, summarized in Fig. 6b. Protocols (ii), (iv), and (v) achieved comparable inference accuracy, while protocol (i) performed the worst. This is expected, since for small *z* values, the GPGF is dominated by terms *z*^*n*^ with *n <* 0, causing the optimization to overemphasize the tails of the sRNA distribution and degrade inference accuracy. In Fig. 6c, we compare the computation times of the three best-performing protocols. Remarkably, protocol (ii) is approximately 8,000 times faster than the traditional MLE+FSP approach and remains faster than protocol (iv). Together, these results demonstrate that the GPGF-based inference method with the integration interval [0.9, 1] achieves the best balance between accuracy and computational efficiency.

## VIII. DISCUSSION

In this work, we investigated the stochastic interaction between mRNA and sRNA molecules. Under the strong antagonistic assumption, the joint probability distribution of mRNA and sRNA is restricted to states in which the two species cannot coexist. While this assumption greatly simplifies the stochastic dynamics, traditional PGF approaches^41^ cannot, in general, yield analytical solutions for such systems, particularly for time-dependent cases even under this simplifying condition. To address this limitation, we introduced a GPGF method, which extends the conventional PGF definition from non-negative counts to include both positive and negative indices. This generalization enables direct analytical treatment of the full stochastic system in the GPGF space, without the need to first derive reduced CMEs under the strong antagonistic assumption. Consequently, the analysis becomes considerably streamlined. The GPGF-based analytical solution can be obtained by identifying the appropriate kernels and applying standard partial differential equation solution techniques. Notably, the resulting expressions typically involve only elementary functions, avoiding hypergeometric terms and thus enhancing numerical stability during evaluation. The corresponding joint distribution can then be recovered by performing a Laurent series expansion of the GPGF, which can be efficiently computed using the NSeries command in Mathematica. The availability of analytical GPGF solutions also facilitates rapid parameter inference from experimental data. Compared with conventional MLE+FSP method, our approach accelerates parameter estimation by approximately three orders of magnitude. This improvement opens the possibility of inferring gene–gene interactions from single-cell RNA sequencing data at high throughput within practical computational times.

Despite its advantages, the GPGF approach also introduces certain limitations. In particular, it does not provide closed-form analytical expressions for statistical moments. In the traditional PGF framework, for example, the mean of a species can be obtained by differentiating the PGF with respect to *z* and evaluating at *z* = 1. For the GPGF, however, this operation yields only the mean of the difference between mRNA and sRNA counts, rather than their individual means. The same limitation extends to higher-order moments. Consequently, moments in our framework must be computed numerically by first expanding the GPGF solution via Laurent series to obtain the joint distribution, and then calculating the desired moments from this distribution.

The four models studied here (Models I–IV) describe mRNA and sRNA degradation as first-order reactions, consistent with the assumption that degradation dominates over dilution caused by cell division. This assumption is well supported for a broad range of genes^43^. However, it may not hold for protein dynamics, where degradation is often slower than dilution^**?**^. Extending the current framework to such cases presents an intriguing direction for future work. For long-lived proteins, one can employ the recent approach proposed in Ref.^50^ to derive the joint distribution of mRNA, sRNA, and protein, thereby obtaining a more comprehensive view of gene regulation under mRNA-sRNA interaction.

## ACKNOWLEDGMENT

This work is supported by NSFC Grant (62573195), Shanghai Action Plan for Technological Innovation Grant (23S41900500), and the Natural Science and Engineering Research Council of Canada’s (NSERC’s) Discovery Grant (RGPIN-2024-06015). We thank Dr. Ramon Grima at the University of Edinburgh for stimulating discussions.

## Supplementary Note 1

Here we present the theoretical details for derivign the reduced chemical master equation (CME) in Eq. (5). Let *q*_*i*→*j*_ denote the rate of transition from state *i* to state *j* (also known as the propensity), where *i* and *j* represent arbitrary probabilistic states such as (*n*_m_, *n*_s_). The total leaving rate for state *i* is given by

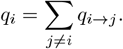

According to Eq. (4), the total leaving rate for state (*m, n*) is

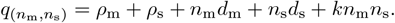

It is observed that

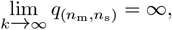

whenever *n*_m_ *>* 0 and *n*_s_ *>* 0. This implies that all probability states satisfying *n*_m_ ≥ 1 and *n*_s_ ≥ 1 are *fast states*. Once the system reaches such a state, it immediately transitions to a neighboring *slow state*, such as (*n*_m_, 0), (0, *n*_s_), or (0, 0) for any *n*_m_ *>* 0 and *n*_s_ *>* 0. We denote the sets of fast and slow states as *A* and *B*, respectively.

Transitions between slow states in *B* can occur in two ways: (i) direct transitions between slow states (*B → B*), and (ii) indirect transitions through a sequence of fast states (*B*→ *A* →· · · → *A* → *B*). The transition rate between any two slow states *i* and *j* can thus be decomposed as

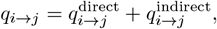

where *i, j* ∈ *B*. The direct transition rate is defined as the product of the total leaving rate from state *i* and the probability of hopping from *i* to *j*, denoted *w*_*i*→*j*_:

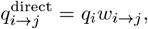

for *i, j* ∈ *B*. The indirect transition rate accounts for all possible paths *c* connecting states *i* and *j* through fast states, and is expressed as

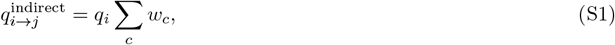

where *w*_*c*_ represents the probability of a specific path *c* between *i* and *j* through fast states,

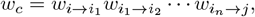

for *i*_1_, *i*_2_, …, *i*_*n*_ ∈ *B*. For large *k*, the transition probability 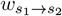 between any two probabilistic states *s*_1_ and *s*_2_ can be computed as

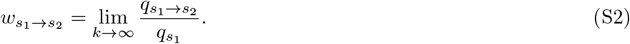

We now use this framework to derive the reduced CME in Eq. (5). Consider transitions of mRNA from one slow state (*n*_m_, 0) to another (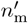, 0). Two cases arise depending on the relationship between *n*_m_ and 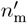.

### Case I (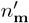*> n*_m_)

This corresponds to mRNA production. There exists only one direct path between (*n*_m_, 0) and (*n*^*′*^m, 0), with 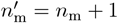. Therefore,

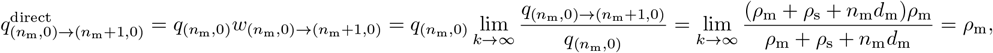

using Eq. (S2). Since no indirect path exists in this case, 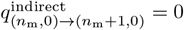, and

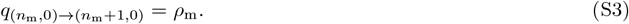

### Case II (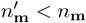)

For 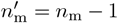, a direct transition exists between (*n*_m_, 0) and (*n*_m_ − 1, 0), with rate

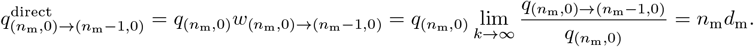

An additional indirect path 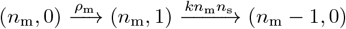 also exists. Using Eq. (S2), the corresponding transition probabilities are

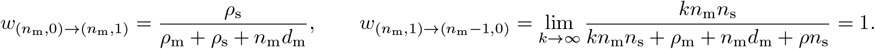

Therefore, the indirect transition rate is

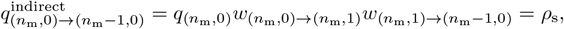

as given by Eq. (S1). Hence, the total transition rate from (*n*_m_, 0) to (*n*_m_ − 1, 0) is

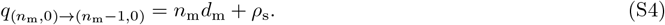

Combining Eqs. (S3) and (S4) yields the first equation in Eq. (5). Owing to the symmetry between mRNA and sRNA in Model I (Fig. 1), the second equation in Eq. (5) follows immediately. Finally, using the first two equations, the reduced CME for the state (0, 0) (the third equation in Eq. (5)) can also be obtained.

## Supplementary Note 2

This section presents the exact solution to the reduced CMEs Eq. (5) using the probability generating function (PGF) method. We first define the generating functions *G*_1_(*z*_1_) and *G*_2_(*z*_2_) for the distributions of mRNA and sRNA:

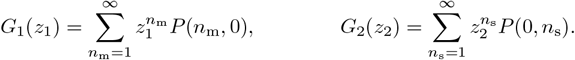

According to this definition, multiplying 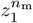 on both sides of the first equation in Eq. (5) and summing from *n*_m_ = 1 to *n*_m_ = +∞, we obtain

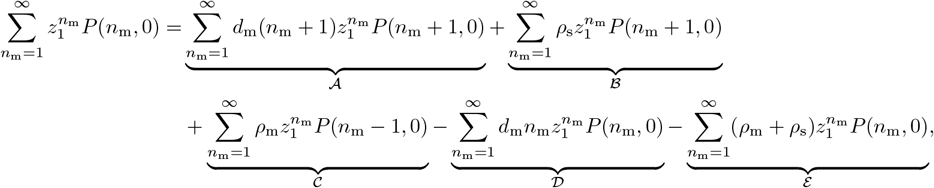

Where

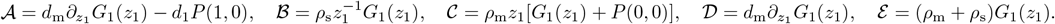

After performing algebraic simplification and applying the steady-state assumption, we obtain

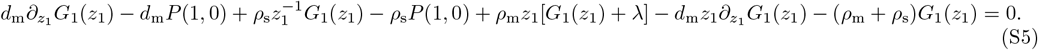

where *λ* := *P* (0, 0). By exploiting the symmetry between mRNA and sRNA, the generating function for the second equation in Eq. (5) becomes

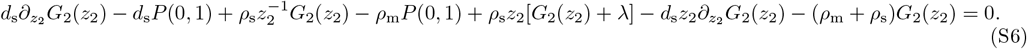

Summing Eqs. (S5), (S6), and the third equation in Eq. (5), we obtain

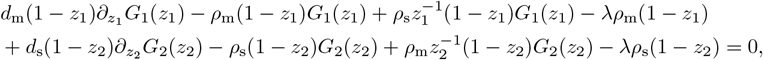

which further implies

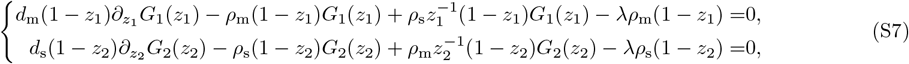

as *z*_1_ and *z*_2_ are independent variables. With initial conditions *G*_1_(0) = 0 and *G*_2_(0) = 0, the solutions to Eq. (S7) are

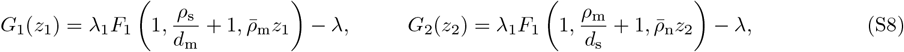

Where 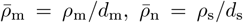, and _1_*F*_1_ denotes the Kummer function. The constant *λ* is determined by the normalization condition *G*_1_(1) + *G*_2_(1) + *λ* = 1, yielding

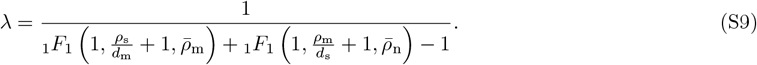

Because mRNA and sRNA cannot coexist at steady state, the generating function can be expressed as

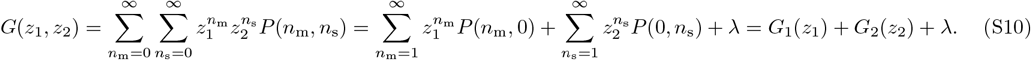

Combining Eqs. (S8), (S9), and (S10) gives Eq. (7). The marginal distributions of mRNA and sRNA are then obtained from

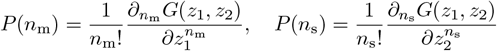

for any *n*_m_ and *n*_s_.

## Supplementary Note 3

### PGF-Based Inference Method

We consider a population of *n*_*c*_ single cells, where the *i*-th cell contains *n*_m,*i*_ mRNA molecules and *n*_s,*i*_ sRNA molecules. The empirical PGF is defined as

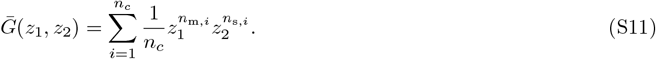

To infer the parameter set *ϕ*, we minimize the discrepancy between the model-predicted PGF, *G*(*z, ϕ*), obtained from Eq. (7), and the empirical PGF, 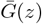. The objective function is defined as

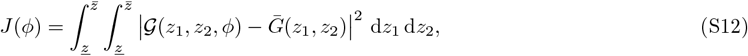

where 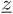 and 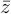 denote the lower and upper bounds of integration, respectively.

To enhance computational efficiency in evaluating Eq. (S12), we approximate the double integral using Gauss–Legendre quadrature:

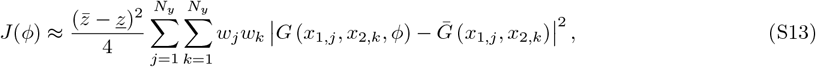

where the quadrature nodes are transformed via

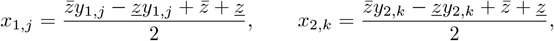

with *y*_*j*_ ∈ [−1, 1] denoting the *j*-th Gauss–Legendre quadrature node of order *N*_*y*_, and *w*_*j*_ the corresponding weight, both computed using the `gausslegendre` function in Julia. All remaining optimization steps follow those described in Algorithm 1, and the complete procedure is summarized in Algorithm S1.

#### Algorithm S1

PGF-based inference method

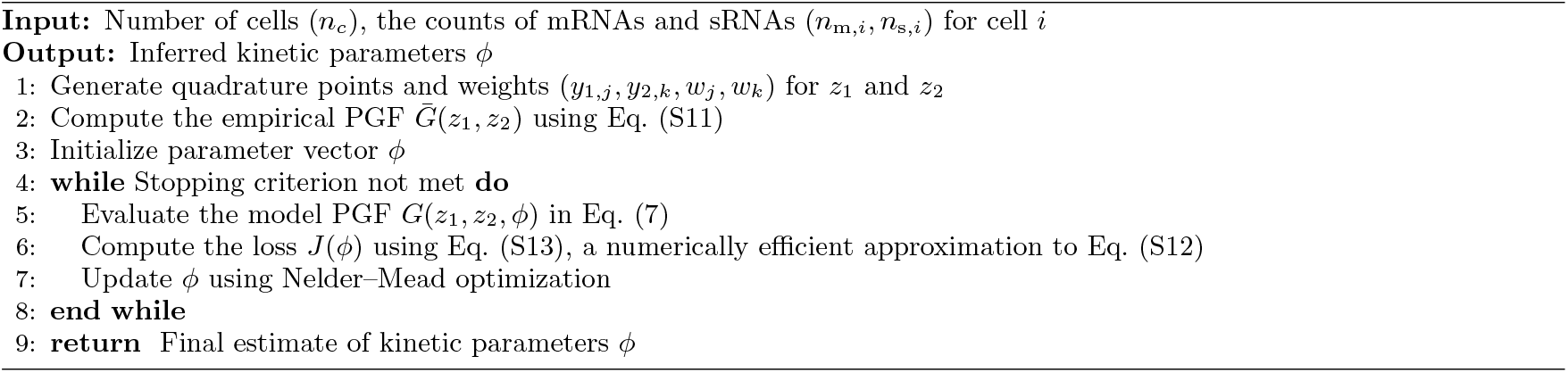

### Maximum Likelihood Estimation (MLE) with the Finite State Projection (FSP) Method

The kinetic parameters *ϕ* are inferred by minimizing the negative log-likelihood function:

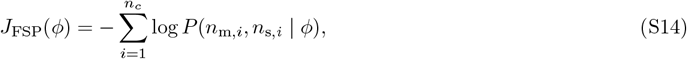

where *P* (*n*_m,*i*_, *n*_s,*i*_ |*ϕ*) denotes the probability of observing *n*_m,*i*_ mRNAs and *n*_s,*i*_ sRNAs in cell *i*, computed using the FSP method to solve Eq. (4).

#### Algorithm S3

MLE Integrated with FSP

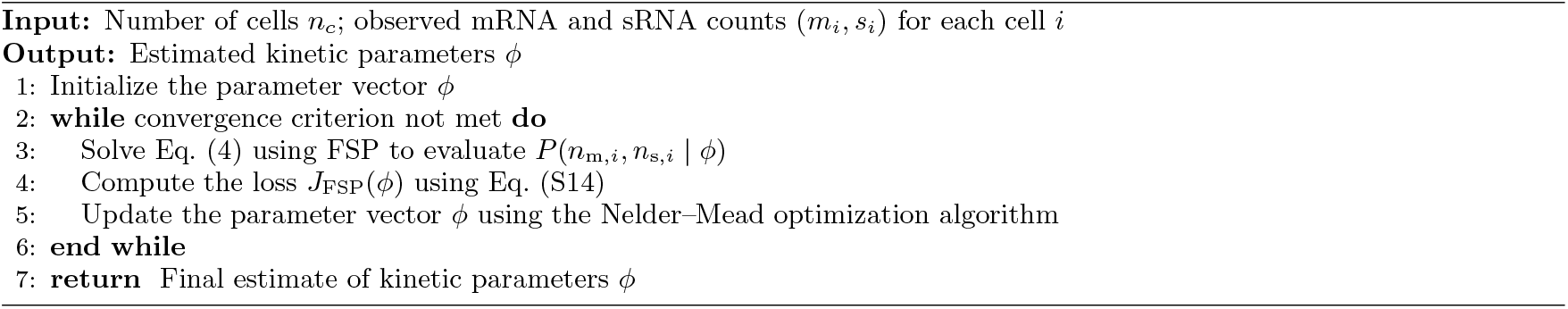

**TABLE S1.** Kinetic parameter sets used to generate synthetic count data for model selection in Fig. 6. Parameters were sampled from the ranges *ρ*_m_ ∈ [4, 40] and *ρ*_n_ ∈ [1, 10], with *n*_*c*_ = 10,000 single cells simulated per parameter set.

|  | Parameters |  |
| --- | --- | --- |
| | $\rho_m$ | $\rho_s$ |
| Set 1 | 10 | 2.5 |
| Set 2 | 15.385 | 4.476 |
| Set 3 | 12.077 | 3.374 |
| Set 4 | 39.261 | 6.424 |
| Set 5 | 13.069 | 2.982 |
| Set 6 | 20.546 | 3.75 |
| Set 7 | 38.691 | 5.226 |
| Set 8 | 4.763 | 1.167 |
| Set 9 | 16.838 | 6.452 |
| Set 10 | 25.878 | 4.903 |

## Notes

### Competing Interest Statement

The authors have declared no competing interest.

